# Emergence of hypervirulent *Klebsiella pneumoniae* ST111 with a novel virulence plasmid causing pyogenic liver abscess

**DOI:** 10.1101/2025.11.25.690365

**Authors:** Liya Feng, Ke Liu, Yuheng Liu, Yanglu Ou, Xiutao Dong, Zhongguo Sun, Liang Chen, Xiaohong Shi, Mingju Hao

## Abstract

Hypervirulent *Klebsiella pneumoniae* (hvKp)–associated pyogenic liver abscess is typically linked to K1/K2 capsule serotypes and canonical lineages such as ST23, ST65, and ST86. Cases outside these lineages are rare but pose significant public health concerns. We investigated an unusual liver abscess in an immunocompetent patient caused by hvKp sequence type 111 (ST111). Five isolates (KP1–KP5) were recovered, all belonging to the same clonal lineage. Genomic analysis revealed a 181 kb IncHI1B/IncFIB virulence plasmid encoding *rmpADC* and the salmochelin cluster (*iroBCDN*), but lacking aerobactin (*iuc*) and *rmpA2* gene loci. Notably, this plasmid also carried the yersiniabactin locus (*ybt*), which is typically located on the chromosome. Comparative genomics indicated that similar ST111 strains have been reported only in China, suggesting clonal expansion of a previously unrecognized hvKp lineage. Functional assays demonstrated that plasmid curing abolished the hypermucoviscous phenotype, reduced capsule production, and completely attenuated virulence in murine models, confirming the plasmid as the major driver of hypervirulence. Interestingly, plasmid loss enhanced biofilm formation and relieved growth burden, highlighting fitness trade-offs associated with virulence. This study documents a rare case of hvKp ST111 liver abscess and identifies a novel plasmid conferring hypervirulence. Our findings expand the known diversity of hvKp and underscore that non-canonical lineages can acquire potent virulence determinants enabling severe disease. Continued clinical and genomic surveillance is essential to detect emerging hvKp clones before widespread dissemination.

## INTRODUCTION

Over the past three decades, hypervirulent *K. pneumoniae* (hvKp) has emerged globally as a leading cause of pyogenic liver abscess (PLA), with a high prevalence in East and Southeast Asia (1, 2). Unlike classical *K. pneumoniae* (cKp), hvKp is notable for its ability to cause severe, tissue-invasive disease in otherwise healthy individuals, often accompanied by metastatic complications such as endophthalmitis and meningitis (1, 3, 4).

The hypervirulent phenotype is largely driven by mobile virulence plasmids that encode siderophores (*ybt*, *iuc*, *iro*), capsule regulators (*rmpA*, *rmpA2*), and accessory genes such as *peg344* (5, 6). These factors collectively enhance iron acquisition, capsule production, hypermucoviscosity, and resistance to host immunity, thereby enabling hvKp to establish invasive infections, including PLA.

HvKp has historically been dominated by ST23, ST65 and ST86, particularly strains with the K1 or K2 capsule serotypes, which are widely reported across Asia, Europe, and the Americas (7–9). In China, ST23 remains the principal cause of hvKp liver abscesses, although other hypervirulent lineages have also been documented (10–12). The hvKp lineages typically harbor conserved virulence plasmids such as KpVP-1 and KpVP-2, which encode the *rmpADC* operon and multiple siderophore systems that collectively support mucoviscosity and invasive potential (13, 14).

Recently, sequence type ST111 has been reported in association with hypervirulence, causing bloodstream infections (15). However, ST111 hvKp has not previously been implicated in PLA. In this study, we describe the identification of an hvKp strain belonging to the ST111 lineage in an immunocompetent case of pyogenic liver abscess. This finding expands the known genetic landscape of hvKp and emphasizes the need for continued genomic and epidemiological monitoring to detect emerging hypervirulent variants.

## MATERIALS AND METHODS

### Clinical Data Collection

Clinical information was obtained from the medical records of a patient diagnosed with pyogenic liver abscess caused by hvKp in 2025. Data included demographic characteristics, presenting symptoms, laboratory findings, imaging results, antimicrobial therapy, and clinical outcomes. All information was collected as part of routine hospital procedures and anonymized prior to analysis. HvKp strains used in this study (KP1–KP5) were clinical isolates obtained from the patient. KP1 was recovered from a blood sample, whereas KP2–KP5 were isolated from drainage fluid of the liver abscess.

### Antimicrobial Susceptibility Testing (AST)

Antibiotic susceptibility was assessed using the VITEK® 2 automated system (bioMérieux), following CLSI M100 guidelines. The panel included: ampicillin, piperacillin–tazobactam, cefazolin, cefuroxime, cefepime, aztreonam, imipenem, meropenem, ceftazidime, ceftriaxone, amikacin, tobramycin, gentamicin, ciprofloxacin, levofloxacin, trimethoprim–sulfamethoxazole and colistin.

### DNA Sequencing and Bioinformatics Analysis

Whole-genome sequencing of *K. pneumoniae* JNQH1480 (KP2) was performed using Illumina NextSeq and Oxford Nanopore MinION platforms. Hybrid de novo assembly was carried out with Unicycler v0.5.0 (16), and genome annotation was completed using Bakta v1.4.0 (17) with manual curation.

Strain characterization, including MLST, K-locus, and O-locus typing, was conducted using Kleborate software v3.1.2 (18). Antibiotic resistance genes and plasmid replicons were identified with ABRicate v1.0.1 using the CARD (19) and PlasmidFinder databases (20). Virulence loci, including siderophore systems such as yersiniabactin and aerobactin, were detected using Kleborate and the VFDB (21).

To assess genetic diversity and phylogeny, all *K. pneumoniae* genomes (n=25,896) available in the NCBI RefSeq database as of August 19, 2025 were downloaded and screened for ST111 isolates. SNPs were identified using Snippy v4.4.3 (https://github.com/tseemann/snippy), with recombination regions filtered by Gubbins v2.4.1 (22) and core SNPs extracted via SNP-sites v2.5.1 (23). A phylogenetic tree was constructed using FastTree (24) and visualized with the ggtree R package (25).

Conjugation-related elements were analyzed using OriT Finder2 (26). Comparative analysis of virulence plasmids was performed using BLASTn and visualized with Easyfig (27) or BRIG (28).

### Plasmid Curing Using the CRISPR/Cas9-Based pCasCure System

To eliminate the virulence plasmid from the host strain (JNQH1480), we employed a CRISPR/Cas9-based plasmid curing strategy previously developed in our lab (29).

Single-guide RNAs (sgRNAs) targeting plasmid replicons (N20_HI1B_:TCAAAAACTCAACAGTAGAA) were designed using Geneious Prime (30). To improve cloning efficiency, the original pCasCure vector was modified (31) to include: (1) reverse-oriented BsaI sites between the J23119 promoter and sgRNA scaffold for Golden Gate assembly of N20 sequences; (2) an oriT element for conjugative transfer; and (3) a GFP marker for visual selection. The resulting construct was designated pCasCure-1 (Figure S1).

Target-specific pCasCure-1 plasmids were assembled via Golden Gate cloning and introduced into *E. coli* WM3064. Conjugation into recipient *K. pneumoniae* strains was performed, and transformants were selected on LB agar with 50 µg/mL apramycin. GFP-positive colonies were verified by PCR using an N20-specific forward primer and a universal reverse primer (N20-188R).

To induce CRISPR/Cas9 activity, confirmed transformants were cultured in LB with apramycin and 0.1% arabinose. Cultures were serially diluted and plated on apramycin-containing agar to select for plasmid cured colonies. Successful plasmid curing was confirmed by PCR targeting the replicon HI1B. To remove the pCasCure vector itself, colonies were streaked onto LB agar with 5% sucrose and incubated at 37 ℃ overnight.

### Pulsed-Field Gel Electrophoresis (PFGE)

To assess the clonal relatedness of isolates KP1–KP5, PFGE was performed. In addition, S1 nuclease–PFGE was employed to confirm complete removal of the virulence plasmid from the ST111 *K. pneumoniae* strain following CRISPR/Cas9-mediated curing. Genomic DNA from both wild-type and plasmid-cured strains was embedded in 1% SeaKem Gold agarose plugs and digested with proteinase K. The plugs were subsequently treated with either XbaI or S1 nuclease to linearize circular DNA. DNA fragments were separated by PFGE using a CHEF Mapper XA system (Bio-Rad) under the following conditions: 14 °C, 6 V/cm for 17 hours, with a pulse time ranging from 2.16 to 63.8 seconds. Gels were stained and visualized using a Gel Doc 2000 imaging system.

### Hypermucoviscosity Assay

To assess mucoviscosity, both string test and sedimentation assay were performed. For the string test (32), bacterial colonies grown on sheep blood agar were gently stretched using an inoculation loop; formation of a viscous string ≥5 mm was considered positive. For sedimentation (33), overnight cultures in LB broth (5 mL, 37 ℃) were measured for OD600, then centrifuged at 2,500 × g for 5 minutes. The OD600 of the upper 1 mL supernatant was recorded, and mucoviscosity was expressed as the ratio of supernatant OD600 to the original culture OD600.

### Capsule Staining

Capsule visualization was performed via negative staining with Indian ink followed by crystal violet counterstaining (34). A loopful of overnight culture was mixed with a drop of Indian ink on a clean slide, spread under a coverslip, and air-dried. The slide was then flooded with 0.1% crystal violet for 10 seconds, rinsed gently with distilled water, and air-dried again. Capsules were observed and imaged under a light microscope at 4000× magnification.

### Growth Curve Analysis

To evaluate the fitness impact of the pVir plasmid, bacterial growth was monitored in Mueller-Hinton broth (10). Overnight cultures were washed twice with PBS, adjusted to OD600 = 0.05, and inoculated into 96-well microtiter plates (200 μL per well). Plates were incubated at 37 ℃, and OD600 readings were recorded at regular intervals over 24 hours using a Multiskan™ FC Microplate Photometer (Thermo Scientific). Measurements were collected at defined time intervals and plotted as mean ± standard deviation (SD) from three biological replicates.

### Capsule Isolation and Quantification

Capsular polysaccharide (CPS) production was quantified by measuring uronic acid content (33, 35). Overnight cultures (500 μL) were mixed with 100 μL of 1% Zwittergent 3-14 in 100 mM citric acid buffer (pH 2.0), incubated at 50 ℃ with shaking (170 rpm) for 30 minutes, and centrifuged at 16,000 × g for 2 minutes. Supernatants (300 μL) were transferred to new tubes, and CPS was precipitated with 1.2 mL absolute ethanol on ice for 30 minutes. Pellets were centrifuged at 16,100 × g for 10 minutes at 4 ℃, air-dried, and resuspended in 100 μL distilled water for overnight incubation.

For quantification, 20 μL of CPS solution was mixed with 120 μL of 12.5 mM sodium tetraborate in concentrated sulfuric acid, heated at 100 ℃ for 5 minutes, cooled, and reacted with 2 μL of 0.15% 3-phenylphenol in 0.5% NaOH. After equilibration, absorbance at 520 nm was measured. CPS concentration was calculated using a galacturonic acid standard curve and normalized to the original culture OD600. All measurements were performed in triplicate.

### Biofilm Formation Assay

Biofilm formation was quantified using crystal violet staining (10). Overnight cultures were diluted 1:1000 in LB medium, and 200 μL was added to each well of a U-bottom 96-well plate. After 24-hour incubation at 37 ℃, wells were washed four times with distilled water to remove planktonic cells. Adherent biofilms were stained with 125 μL of 0.1% crystal violet for 10 minutes, washed six times, and solubilized with 150 μL of 30% glacial acetic acid. OD590 was measured using a microplate reader (Thermo Scientific). Each strain was tested in triplicate, and experiments were repeated twice.

### Murine Infection Model

To evaluate the virulence of the ST111-type hvKp strain and assess the contribution of its virulence plasmid, a murine infection model was established using 4-week-old female specific pathogen-free (SPF) BALB/c mice (purchased from Beijing Vital River Laboratory Animal Technology Co., Ltd.).

Mice were randomly assigned to two groups: the experimental group, inoculated with the wild-type hvKp strain, and the control group, inoculated with the plasmid-cured derivative. Each group was further subdivided into four dose cohorts (n = 10 per cohort), receiving intraperitoneal injections of 1×10^4^, 1×10⁵, 1×10⁶, or 1×10⁷ colony-forming units (CFU) per mouse.

Following bacterial challenge, animals were monitored closely for 14 days to assess survival and clinical signs of severe illness. Humane endpoints were defined in accordance with the American Veterinary Medical Association Guidelines, and mice exhibiting any of the following signs were euthanized immediately: hunched posture, ruffled fur, labored breathing, reluctance to move, photophobia, or dehydration. Survival and mortality were recorded as dichotomous outcomes.

### Statistical Analysis

The difference in BALB/c murine survival between groups was assessed using the Log-rank (Mantel–Cox) test, which evaluates statistical significance across Kaplan–Meier survival curves. Statistical comparisons between growth curves were performed using a two-way analysis of variance (ANOVA). A *P*-value < 0.05 was considered statistically significant. All analyses and visualizations were performed using GraphPad Prism (version 10.1.2)

## RESULTS

### Case Presentation

A 39-year-old woman with no history of diabetes or immunosuppression presented in our hospital in January 2025 with acute onset of high fever (maximum 39.5 ℃), chills, fatigue, nausea, anorexia, chest tightness, palpitations, dizziness, and a mild productive cough with scant white sputum. Despite self-administration of acetaminophen and oseltamivir, her fever recurred within hours, accompanied by worsening malaise and upper abdominal discomfort.

On admission, she was febrile (39.6 ℃), tachycardic (130 bpm), and tachypneic (26 breaths/min). Physical examination revealed right upper quadrant tenderness without other significant cardiopulmonary or abdominal findings. Laboratory evaluation demonstrated marked leukocytosis (20.43 × 10⁹/L), elevated C-reactive protein (272.3 mg/L), and a markedly increased procalcitonin level (22.343 ng/mL) (Figure 1). Abdominal imaging, including CT, ultrasound, and MRI, identified multiple hepatic abscesses in the right lobe. The admission MRI revealed several mass-like lesions characterized by long T1 and T2 signals. The largest lesion measured approximately 10.4 × 7.1 cm and contained multiple cystic and septated regions with elevated T2 signal intensity. Diffusion-weighted imaging showed hyperintensity with reduced ADC values, while contrast-enhanced sequences demonstrated pronounced enhancement of lesion margins and septa (Figure 2).

**Figure 1.**
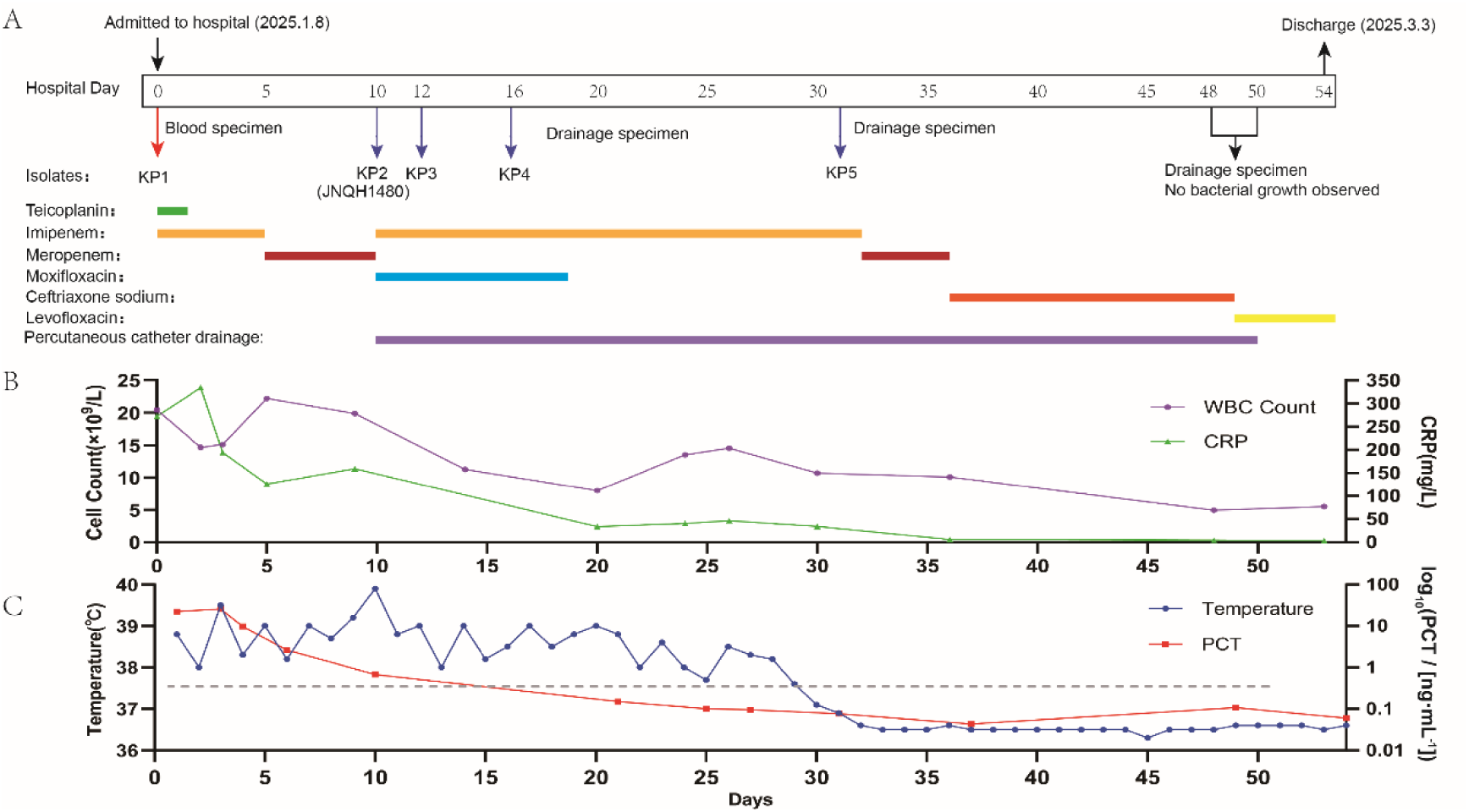
Timeline of antimicrobial therapy, percutaneous catheter drainage, and clinical parameters during hospitalization. (A) Timeline illustrating the isolation of hvKp and the corresponding course of antimicrobial treatment, together with the timing of percutaneous catheter drainage. (B) Trends in leukocyte count, C-reactive protein, procalcitonin, and body temperature throughout the hospitalization period. The x-axis represents days of hospitalization, and the y-axis indicates the respective clinical parameters.

**Figure 2.**
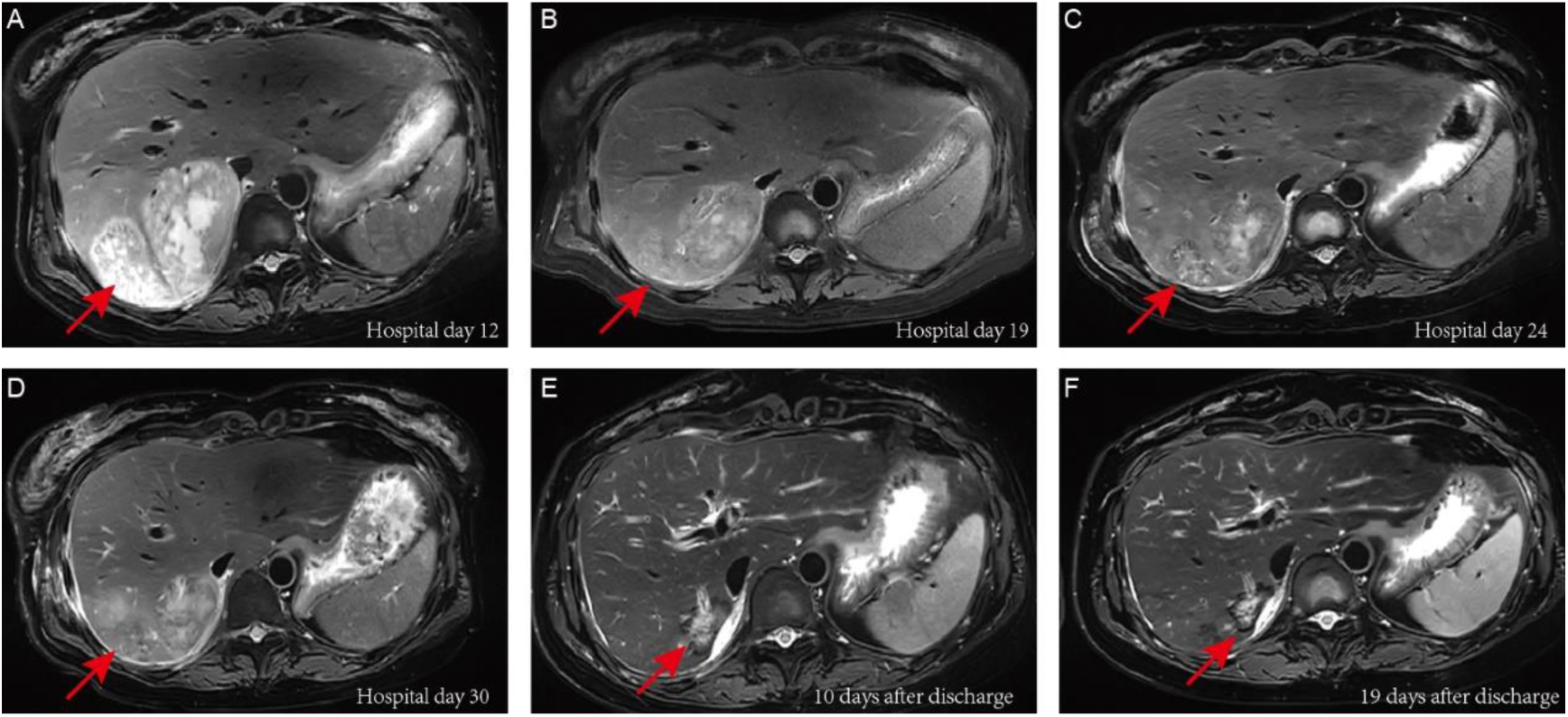
Characteristic MRI findings of hepatic abscesses for the patient. Sequential liver MRI scans demonstrating the evolution and resolution of hepatic abscesses on T2-weighted imaging. (A) The initial MRI at admission revealed multiple mass-like lesions in the right hepatic lobe. (B-D) Post-procedural MRI following antibiotic treatment and percutaneous catheter drainage showed irregular long T2 signals with a marked reduction in lesion size (red arrows). (E-F) Outpatient follow-up MRIs performed after discharge confirmed substantial improvement and near-complete resolution of the hepatic abscesses (red arrows). The corresponding time for each MRI examination is indicated in the images.

Blood cultures obtained prior to empirical antimicrobial therapy with teicoplanin (200 mg, q12h, IV) and imipenem (1 g, q8h, IV) grew *K. pneumoniae* (strain KP1). Antimicrobial susceptibility testing indicated broad susceptibility. On hospital day 10, percutaneous catheter drainage was performed, with repeated cultures from the drainage fluid (strains KP2–KP5) consistently yielding *K. pneumoniae*. Species identification of all isolates was confirmed using MALDI-TOF mass spectrometry. Based on susceptibility results, therapy was escalated to imipenem (0.5 g, q6h, IV) combined with moxifloxacin (0.4 g, qd, IV), followed by imipenem monotherapy.

On hospital day 31, *K. pneumoniae* was still cultured from the drainage specimen; however, the patient’s clinical condition and inflammatory markers showed steady improvement (Figure 1). On hospital day 36, antibiotics were de-escalated to ceftriaxone (2 g, q12h, IV), and subsequently transitioned to levofloxacin (0.5 g, qd, IV). By hospital days 48 and 50, drainage cultures were consecutively negative, and MRI demonstrated marked resolution of the hepatic abscesses. The catheter was removed, and the patient was discharged on hospital day 54. Outpatient follow-up MRIs confirmed substantial improvement and near-complete resolution of the hepatic abscesses, with the patient remaining clinically stable and fully recovered (Figure 2).

### Antimicrobial Susceptibility

All *K. pneumoniae* isolates (KP1–KP5) exhibited identical antimicrobial susceptibility profiles. The strains were uniformly resistant to ampicillin, while remaining broadly susceptible to other major classes, including β-lactams (cephalosporins, carbapenems, and β-lactam/β-lactamase inhibitor combinations), aminoglycosides, fluoroquinolones, and colistin (Table S1).

### Clonal Relatedness and Genomic Features of ST111 hvKp strains

PFGE analysis revealed that all isolates (KP1-KP5) exhibited identical banding patterns (Figure 3), indicating that the strains originated from the same clonal lineage. Genome sequencing and assembly of strain JNQH1480 produced a single 5.02 Mb chromosome. Multilocus sequence typing assigned JNQH1480 to ST111. In silico serotyping identified a KL63 capsular locus and an O1/O2v2 O-antigen locus. In addition to its chromosome, JNQH1480 harbors a 181,651 bp virulence plasmid (pJNQH1480) encoding 188 coding sequences. Replicon analysis classified this plasmid as IncHI1B/IncFIB. Key virulence determinants on the plasmid include the *rmpADC* mucoid-regulating operon, the salmochelin biosynthesis cluster (*iroBCDN*), and the yersiniabactin locus (*ybt* and *irp*). Notably, compared with the canonical pK2044 plasmid, the plasmid lacks a complete *rmpA2* and the *iucABCD-iutA* aerobactin operon. The plasmid only contains *TraI* relaxase genes, lacking other conjugation elements.

**Figure 3.**
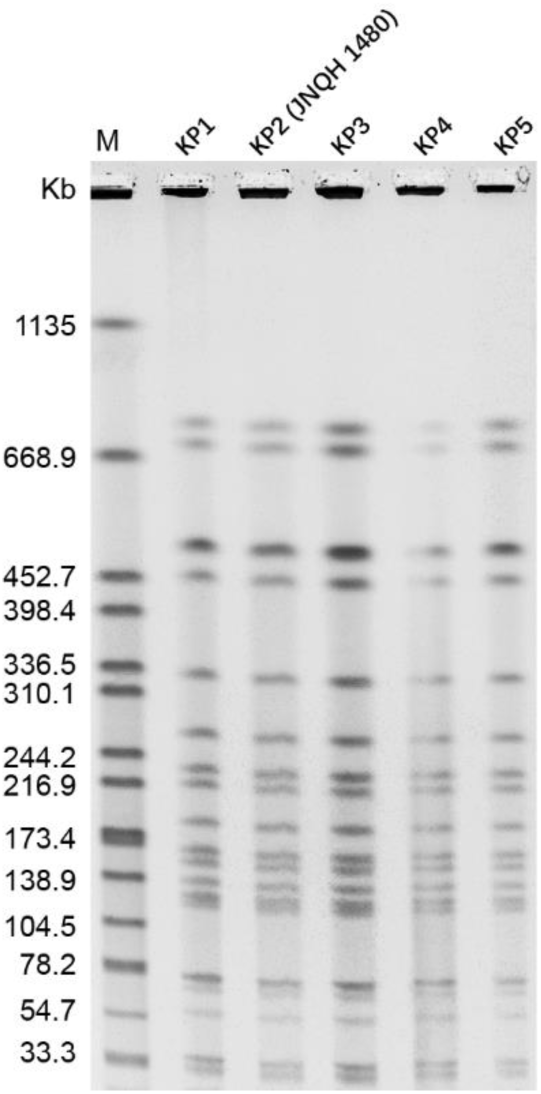
PFGE analysis of hvKp isolates collected from the patient. PFGE profiles demonstrated that all isolates (KP1–KP5) exhibited identical banding patterns. Strain identifiers are indicated above each lane. The molecular size marker (H9812) is shown in lane M, with fragment sizes annotated on the left.

### Epidemiological Distribution and Genomic Characteristics of ST111 hvKp

To investigate the global distribution and genomic features of sequence type 111 (*K. pneumoniae* ST111), we analyzed all publicly available genome assemblies in the NCBI database as of August 19, 2025, identifying a total of 87 ST111 isolates. Phylogenetic analysis (Figure 4) revealed a distinct clade comprising 17 closely related isolates, all originating from China, suggesting a potential regional clonal expansion of an emerging hypervirulent lineage.

**Figure 4.**
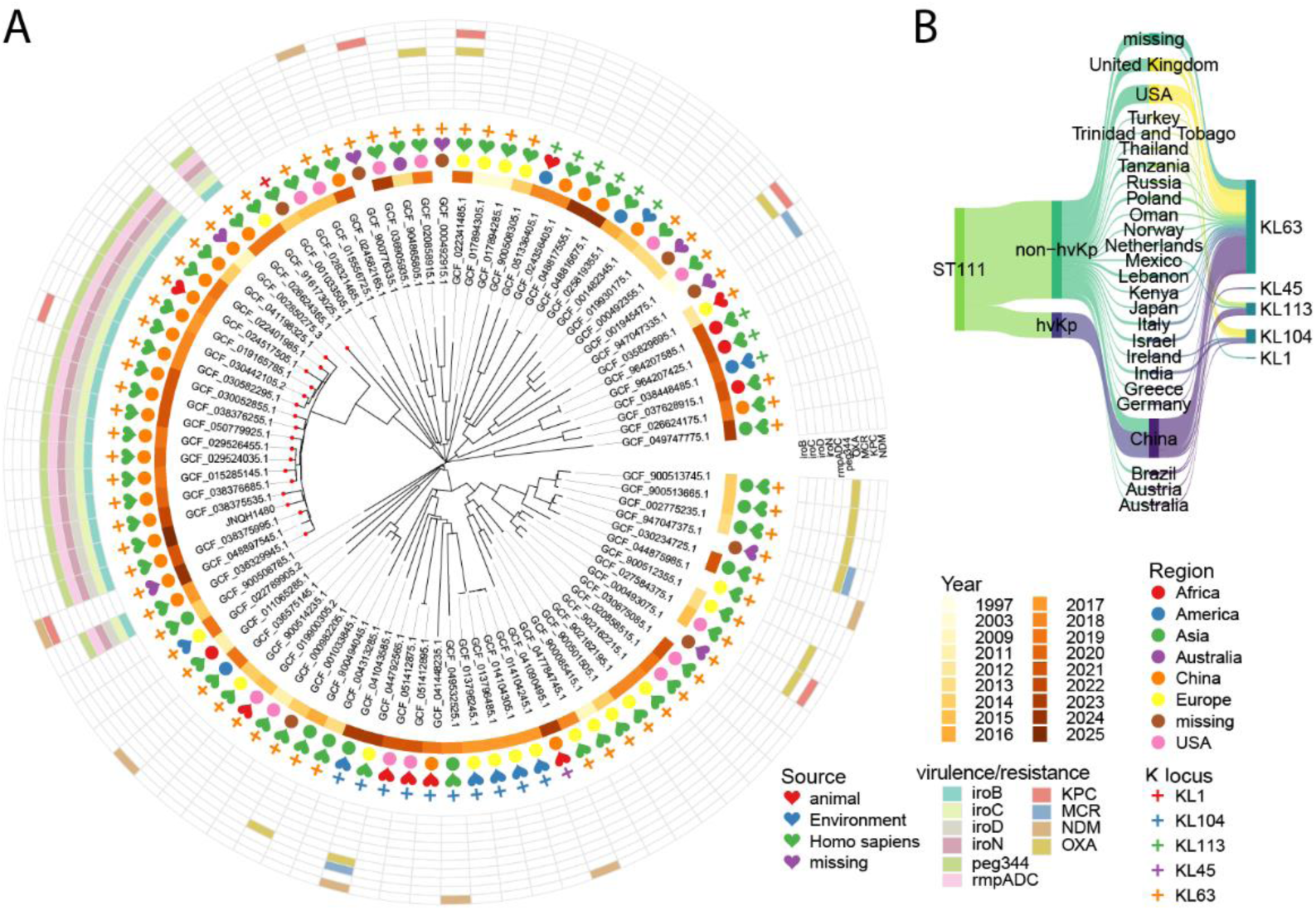
**Phylogenetic relationships and virulence profiles of ST111 *Klebsiella pneumoniae.*** (A) Maximum-likelihood phylogeny of 88 ST111 *K. pneumoniae* isolates, annotated with metadata including collection year, country of origin, isolation source, virulence loci, and resistance genes. Tip nodes corresponding to strains harboring hvKp-associated loci—*iro*, *rmpADC*, and *peg344*—are highlighted in red. (B) All ST111 hvKp strains were isolated in China and uniformly classified as KL63. In contrast, non-hvKp ST111 strains showed global distribution.

All 17 isolates within this clade harbored hallmark hypervirulence determinants, including *rmpADC*, *iro*, and *peg-344*, and uniformly possessed the KL63 capsular locus. Interestinly, these strains were recovered from multiple provinces across China, indicating broad geographic dissemination. The earliest isolate (GCF_041198325.1) was recovered from a lung infection in Zhejiang Province in 2014, while another (GCF_019165785.1) associated with a liver abscess was identified in Hunan Province.

With the exception of one environmental isolate (GCF_024517505), all hypervirulent ST111 strains were derived from human clinical samples (Figure 4A). Notably, only one strain (GCF_030582295.1), isolated from a blood sample in China in 2018, carried the carbapenemase gene *bla*_KPC-2_. The remaining hypervirulent isolates lacked carbapenemase or extended-spectrum β-lactamase (ESBL) genes.

### Comparative Analysis of the Virulence Plasmid pJNQH1480

To characterize the virulence plasmid carried by strain JNQH1480, we performed comparative BLAST analysis, identifying 21 related plasmids with >99.7% sequence identity and >80% coverage (Table S2; Figure 5). These plasmids ranged from 181–360 kb and were predominantly isolated in China between 2013 and 2024. All plasmids belonged to the IncHI1B/IncFIB replicon type and were distributed across diverse *K. pneumoniae* STs and even *E. coli*, supporting their potential for horizontal dissemination. All plasmids retained a complete yersiniabactin (*ybt*) cluster whereas some plasmids lack the canonical virulence associated genes (Figure 5).

**Figure 5.**
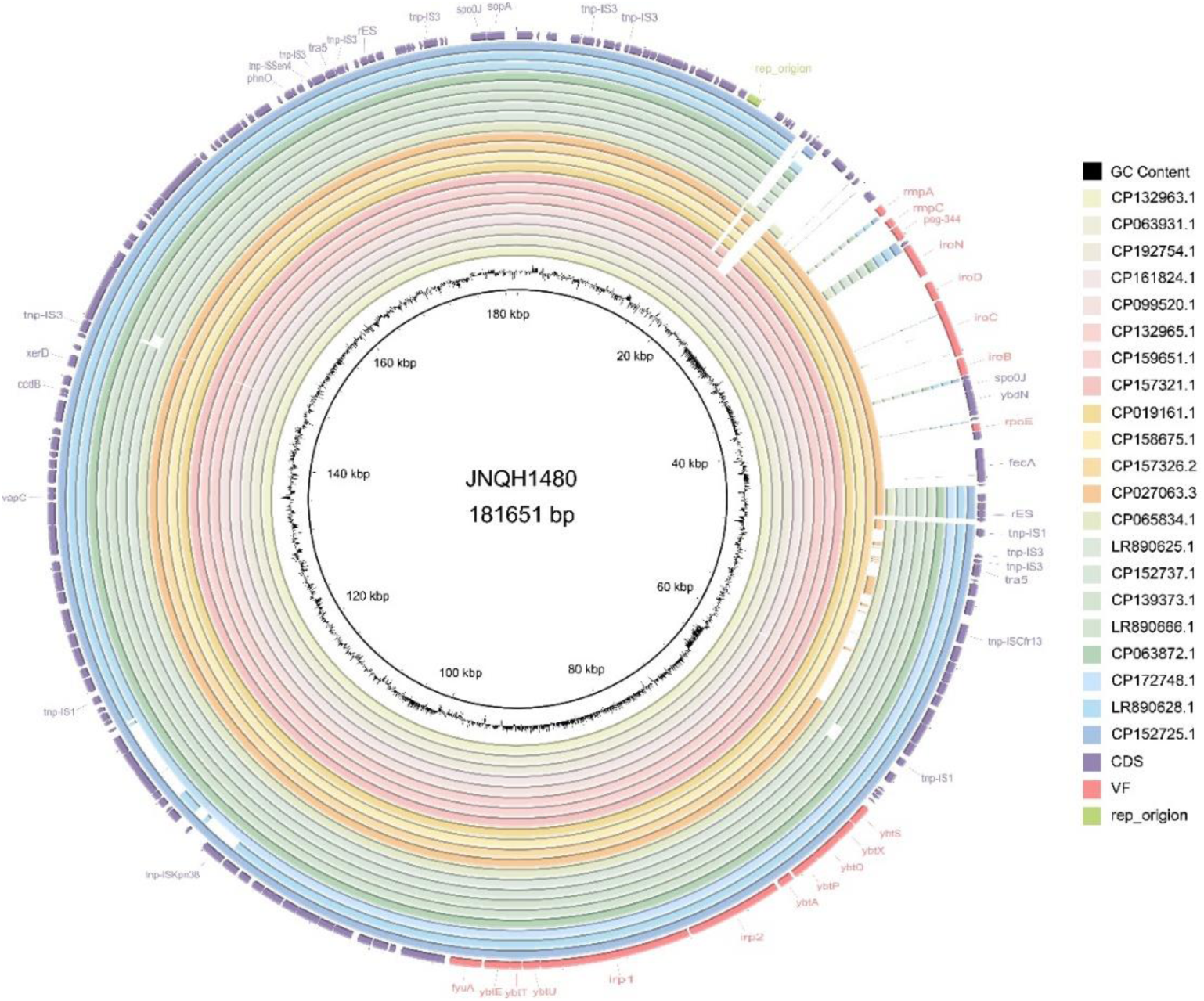
Circular visualization of the virulence plasmid in ST111 hvKp. The circular map displays comparative alignment of virulence plasmids from multiple strains. Concentric rings, arranged from the center outward, represent plasmids with increasing sequence similarity to the reference. The outermost ring corresponds to the plasmid pJNQH1480 characterized in this study, with annotated virulence genes and replicon types highlighted in red.

Based on virulence-gene content and genomic synteny, the plasmids segregated into three patterns (Figure 6A). Pattern 1 plasmids contained the *ybt* cluster, *irp* and *fyuA* virulence genes, and lacked canonical hypervirulence *rmpADC* loci. These plasmids were globally distributed. Pattern 2 plasmids (n = 7) shared highly conserved architecture with the plasmid pJNQH1480 (>99.9% nucleotide identity), despite being distributed across different STs. Four of these plasmids were found in ST111 strains. The pJNQH1480 plasmid was nearly identical to pKP35_vir and pKP36_vir (99.97% identity, 100% coverage), both previously isolated from bloodstream infections in China (15). An inversion relative to pKP36_vir was observed in pJNQH1480, mediated by inverted repeat sequences (Figure 6B). Pattern 3 plasmids carried the *ybt* cluster together with canonical hypervirulence loci (*rmpA, rmpA2, iut, iro*). This group included plasmid pGN-2, a 2015 isolate from China carrying a pK2044-like virulence cluster (36). While pGN-2 and pK2044 shared 99.97% sequence identity across the virulence loci, they exhibited only 50% coverage of the plasmid backbone (Figure 6B). Notably, one plasmid harbored the *bla*_KPC-2_ resistance gene and was identified as a hybrid plasmid comprising HI1B/FIB and IncFII replicon types in *Escherichia coli*. This finding suggests a convergence of hypervirulence and carbapenem resistance, with potential for horizontal transfer among diverse *Enterobacteriaceae* species (7). Interestingly, plasmids in Patterns 2 and 3 were exclusively identified in China, whereas Pattern 1 plasmids displayed broad geographic distribution.

**Figure 6.**
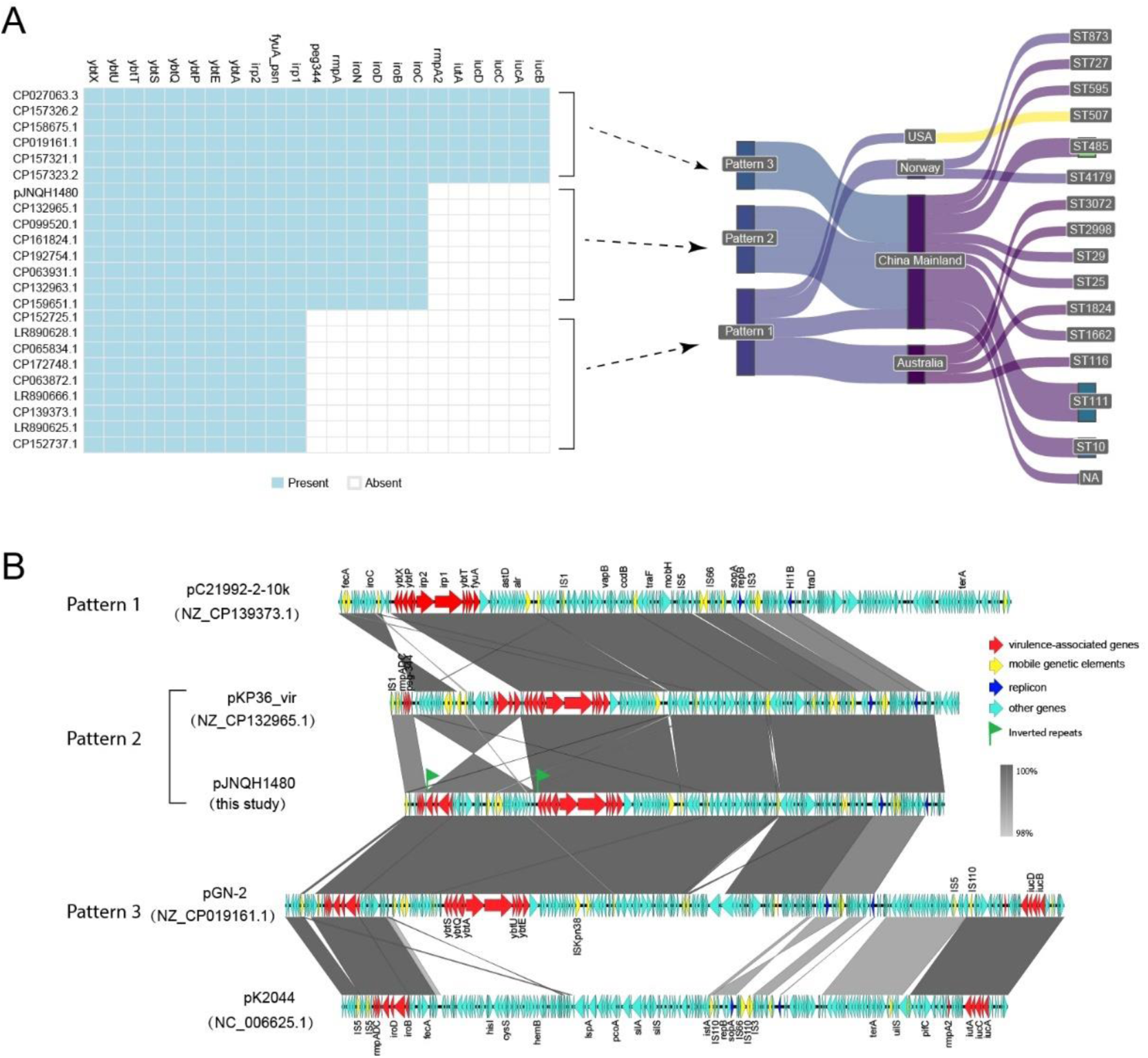
Genomic features, virulence profiles, and distribution of KpVP-3–like plasmid variants. (A) Virulence gene profiling revealed three distinct patterns among KpVP-3–like plasmids. Pattern 1 plasmids encode the *ybt* siderophore cluster along with *irp* and *fyuA*. Pattern 2 plasmids additionally harbor *iro*, *rmpADC*, and *peg344*. Pattern 3 plasmids contain the full canonical virulence repertoire, including *rmpADC*, *rmpA2D2*, *iuc*, *iut*, *peg344*, *iro*, *ybt*, *irp*, and *fyuA*. Geographic and host distribution analysis showed that Pattern 1 plasmids were widespread across diverse regions and sequence types (STs). In contrast, Pattern 2 and Pattern 3 plasmids were exclusively identified in China, restricted to *K. pneumoniae*, but spanning a broad range of STs. (B) Linear alignment of the pJNQH1480 plasmid with representative plasmids in three patterns and the canonical pK2044 (AY378100), highlighting conserved and divergent regions. Red, yellow, dark blue, and light blue arrows indicate virulence genes, mobile genetic elements, replicon types, and other functional regions, respectively.

Collectively, these findings suggest that the pJNQH1480 plasmid likely originated from a pK2044-like ancestor, retaining *rmpA* and *iroN* while losing *rmpA2* and *iuc*, thereby forming a distinct virulence architecture. The consistent presence of *rmpADC* across hypervirulent ST111 isolates underscores its central role in mediating hypervirulence in this lineage.

### Plasmid Curing and Validation of pVir Removal

To assess the phenotypic impact of the virulence plasmid (pVir) on the bacterial host, we employed the CRISPR/Cas9-based pCasCure system (37) to selectively and precisely eliminate pVir from the wild-type strain. Ten colonies were randomly selected for verification, and all (10/10) tested negative for the HI1B replicon (Figure S2), confirming successful plasmid curing. Further validation using S1-pulsed-field gel electrophoresis (S1-PFGE) demonstrated complete loss of the plasmid, as indicated by its absence in the cured strain (Figure 7A).

**Figure 7.**
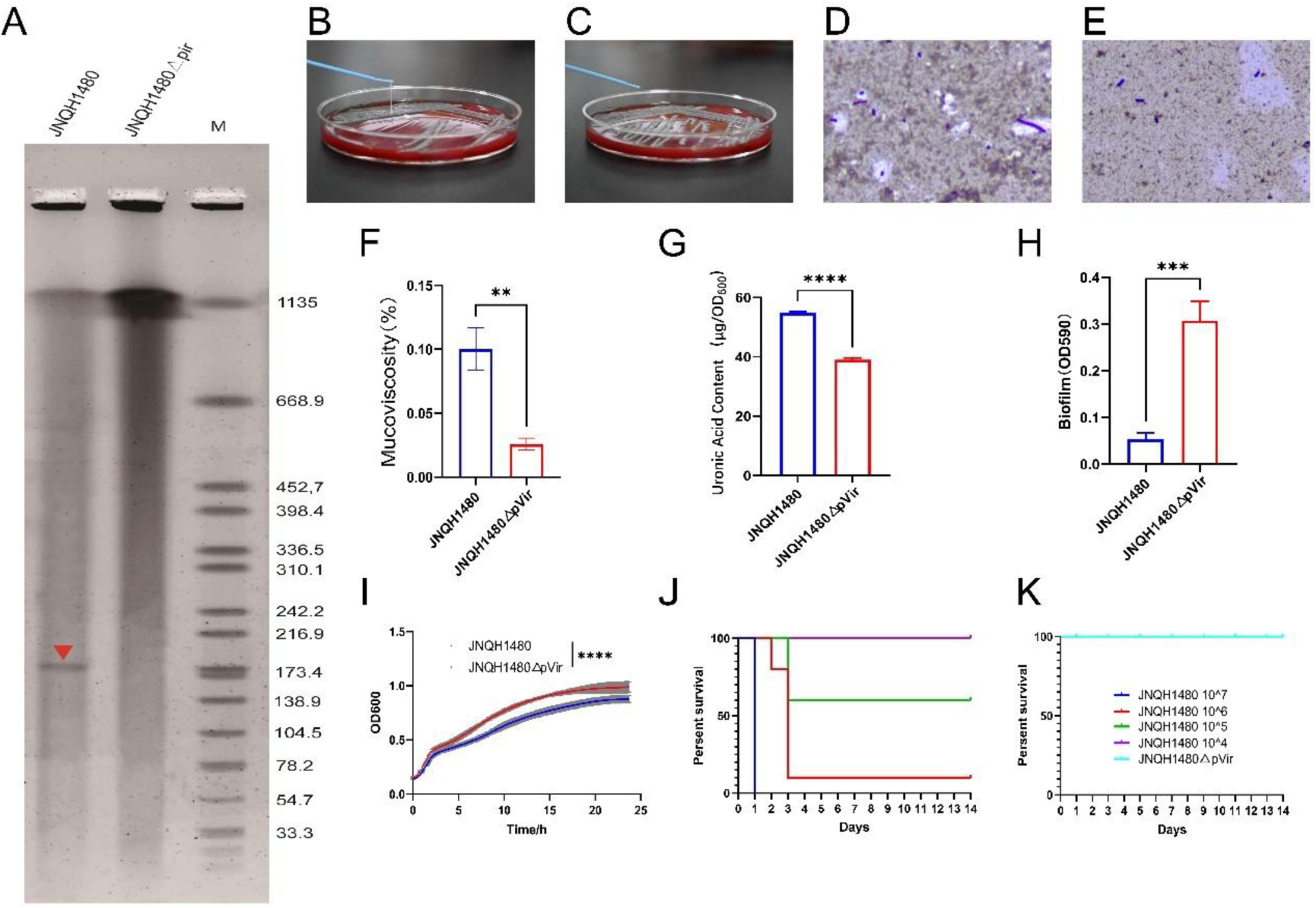
Phenotypic and virulence characterization of ST111 hvKp JNQH1480 and its plasmid-cured derivative (ΔpVir) (A) S1-PFGE analysis confirming complete elimination of the pVir plasmid in the ΔpVir strain. The arrow indicates the pVir band present in the wild-type JNQH1480 strain. M: molecular weight marker. (B–C) String test demonstrating hypermucoviscosity in the wild-type strain (B), which is absent in the plasmid-cured derivative (C). (D–E) India ink capsule staining of wild-type (D) and ΔpVir mutant (E) strains. (F-I) Sedimentation assay (F), growth curves (G), quantification of capsule production (H) and Biofilm formation assay (I) of wild-type and ΔpVir strains. Data are shown as mean ± SD (n = 3 biological replicates; \*\**P*<0.01, *** *P*<0.001, **** *P*<0.0001, unpaired *t*-test). (J-K) Survival curves of mice following intraperitoneal injection with varying doses of wild-type JNQH1480 (J) and ΔpVir strain (K) (n = 10 per group). The calculated LD₅₀ for the wild-type strain was 1.4 × 10⁵ CFU, whereas all mice infected with the pVir-cured strain survived through 14 days.

### Impact of pVir Curing on In Vitro Virulence Phenotypes and Pathogenicity in ST111 HvKp

Elimination of the pVir plasmid from the ST111-hvKp clinical strain JNQH1480 resulted in marked alterations in virulence-associated phenotypes. The wild-type strain displayed pronounced hypermucoviscosity, producing positive string test and exhibiting substantially decreased sedimentation, whereas the pVir-cured mutant lost the hypermucoviscous phenotype, showing a negative string test (Figure 7B-C) and >70% increase in sedimentation (Figure 7F). Capsule visualization confirmed a robust capsule in the wild-type strain, while the cured derivative displayed a visibly reduced capsule layer (Figure 7D-E). Consistently, uronic acid quantification demonstrated a ∼28.6% reduction in capsule production following pVir removal (Figure 7G). Interestingly, biofilm production increased approximately 5.7-fold in the pVir-cured strain compared with the wild type (Figure 7H), indicating that pVir enhances capsule formation while simultaneously suppressing biofilm development. In addition, assessment of bacterial *in vitro* fitness by growth curve revealed that the presence of pVir imposed a significant growth burden, highlighting a cost associated with maintaining the virulence plasmid (Figure 7I). The overall MIC profiles remained largely unchanged after pVir curing (Table S1).

To determine the contribution of pVir to pathogenicity, we performed a murine lethality assay using BALB/c mice challenged intraperitoneally with graded bacterial doses (10⁴–10⁷ CFU). The wild-type strain induced dose-dependent mortality, with 100% lethality observed at 10⁷ CFU within 24 hours, 90% mortality at 10⁶ CFU within 3 days, and 40% mortality at 10⁵ CFU over 14 days, whereas all mice survived through 14 days at 10⁴ CFU (Figure 7J). In contrast, all mice infected with the pVir-cured strain survived even at the highest inoculum (10⁷ CFU) throughout the 14-day observation period (Figure 7K). The calculated LD₅₀ for the wild-type strain was 1.4 × 10⁵ CFU whereas the pVir-cured strain had a LD₅₀ >10⁷ CFU, confirming pVir as a critical determinant of in-vivo virulence.

## DISCUSSION

Pyogenic liver abscesses caused by hvKp represent a major clinical challenge, as these infections are often refractory to antibiotic therapy and carry a substantial risk of dissemination to distant sites such as the eye or central nervous system, leading to vision loss or death (38–40). Reported mortality rates for hvKp liver abscesses range from 5%–40%, underscoring their clinical severity (1). Several microenvironmental conditions known to promote antibiotic tolerance in other bacterial species—including nutrient limitation, oxidative and nitrosative stress, hypoxia, and acidic pH—have been documented within liver abscesses, where restricted antibiotic penetration further contributes to treatment failure (1, 41). Although *Klebsiella* liver abscesses are most commonly associated with K1 and K2 serotypes, the present study identifies and functionally characterizes a novel virulence plasmid in a rarely reported ST111 hvKp isolate. Despite in vitro susceptibility to carbapenems and other agents, the patient experienced a protracted disease course requiring nearly two months of carbapenem therapy and repeated drainage procedures, highlighting the capacity of hvKp to cause persistent, deep-seated infections even in immunocompetent individuals and despite appropriate antimicrobial treatment.

Our genomic analysis demonstrates that ST111 represents an emerging hypervirulent lineage with a distinct evolutionary trajectory. Phylogenetic clustering, KL63 dominance, and geographic concentration in China suggest recent clonal expansion. Notably, the conserved presence of the *rmpADC* operon, despite partial loss of canonical virulence loci (*rmpA2, iut, iuc*), highlights its central role in sustaining hypervirulence. The pVir plasmid in this strain shared <40% homology with known KpVP-1/2 backbones and has recently been designated KpVP-3 (15), expanding the spectrum of recognized hvKp virulence plasmids. Unlike classical hvKp plasmids harboring aerobactin and colibactin clusters, the ST111 plasmid encoded *rmpADC, iroBCDN,* and *ybtAPQSX*, identifying an alternative virulence gene configuration capable of driving severe disease.

These findings refine current biomarker-based definitions of hvKp. While the presence of *iucA, iroB, peg-344, rmpA*, and *rmpA2* is widely used to predict hypervirulence (42–45), our isolate lacked *iucA* and *rmpA2*, yet exhibited a very low LD₅₀ in vivo, indicating clear hypervirulence (46). Prior studies show that yersiniabactin alone does not define hvKp but increases virulence in cKp backgrounds) (46). Thus, the ST111 strain represents an important example in which *rmpADC* and *iroBCDN*, together with *ybt*, are sufficient to confer a hypervirulent phenotype, expanding the known genetic basis of hvKp.

Functional validation through CRISPR-mediated plasmid curing confirmed pVir as the major virulence determinant (37). Loss of pVir eliminated hypermucoviscosity, markedly reduced capsule thickness, and attenuated virulence in a murine model—consistent with the pivotal regulatory role of *rmpA* in capsule biosynthesis (47). Interestingly, plasmid curing enhanced biofilm formation, likely due to reduced capsule masking and increased fimbrial exposure (48), and relieved a growth fitness burden. These findings demonstrated trade-offs between hypervirulence and fitness, wherein plasmid carriage promotes host invasion but incurs metabolic costs that may reduce environmental competitiveness (49, 50).

Our genomic analysis revealed that virulence plasmids with highly conserved gene repertoires are distributed across diverse *K. pneumoniae* lineages. Although the pVir appears to lack conjugation machinery and demonstrates no self-transfer capability in vitro (15), the detection of closely related virulence plasmids in phylogenetically distinct *K. pneumoniae* strains and even *E. coli* underscores the potential for horizontal dissemination. Recent studies have shown that non-conjugative plasmids can be mobilized by co-resident helper plasmids or through the formation of hybrid plasmids that combine virulence and resistance determinants (51–53), raising concerns that pVir-like architectures may continue to spread despite their intrinsic transfer limitations. The phylogenetic clustering of ST111 isolates harboring nearly identical virulence plasmid profiles further supports a dual evolutionary pathway for hypervirulence dissemination: clonal expansion of ST111 and plasmid-mediated horizontal gene transfer. Such a dual dissemination strategy heightens the risk of both the sustained propagation of established hypervirulent lineages and the emergence of novel hvKp variants through plasmid acquisition. Given these dynamics, ongoing genomic surveillance and evolutionary monitoring of virulence plasmids—particularly within emerging ST111 clades—are essential to anticipate and mitigate future hvKp threats.

Taken together, our findings broaden the genetic landscape of hvKp beyond classical lineages, establish ST111 as a clinically significant hypervirulent clone in China. These findings highlight the urgent need for integrated genomic surveillance, improved diagnostic tools, and novel therapeutic approaches targeting virulence mechanisms to prevent the further dissemination of hvKp in both community and healthcare settings.

## ACKNOWLEDGEMENTS

This work was supported by the Shandong Provincial Natural Science Foundation (grant number ZR2021MH078).

## FUNDING

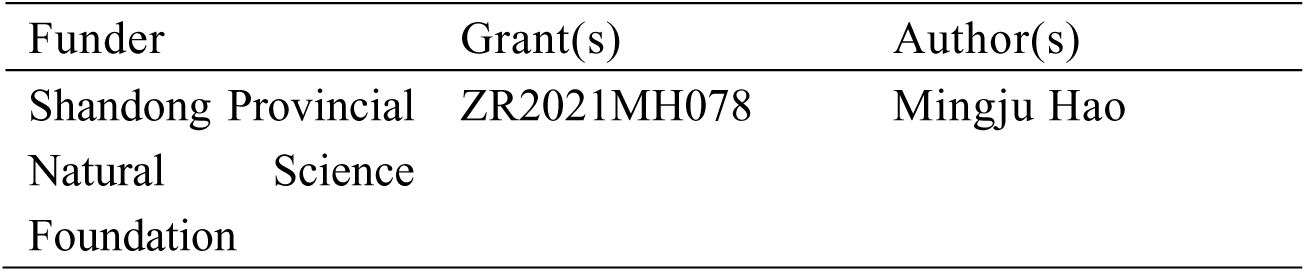

## AUTHOR CONTRIBUTIONS

Liya Feng, Data curation, Methodology, Writing – original draft | Ke Liu, Data curation, Methodology, Writing – original draft | Yuheng Liu, Data curation | Yanglu Ou, Methodology | Xiutao Dong, Resources | Zhongguo Sun, Methodology | Liang Chen, Writing –review and editing | Xiaohong Shi, Project administration, Resources, Supervision | Mingju Hao, Funding acquisition, Methodology, Software, Supervision, Writing – original draft, Writing –review and editing.

## DATA AVAILABILITY

The complete genome sequence of strain JNQH1480 has been deposited at DDBJ/ENA/GenBank under the accession JBRXBC000000000. The version described in this paper is version JBRXBC010000000. The annotated sequence of the pCasCure-1 vector utilized in this study has been deposited in GenBank under accession number PV792961.

## ETHICS APPROVAL

The study was conducted in accordance with the Declaration of Helsinki and approved by the Institutional Review Board (IRB) of the First Affiliated Hospital of Shandong First Medical University & Shandong Provincial Qianfoshan Hospital. In addition, animal experiments were reviewed and approved by the Animal Ethics Committee of the same institution.

## Supplemental Material

**Figure S1.**
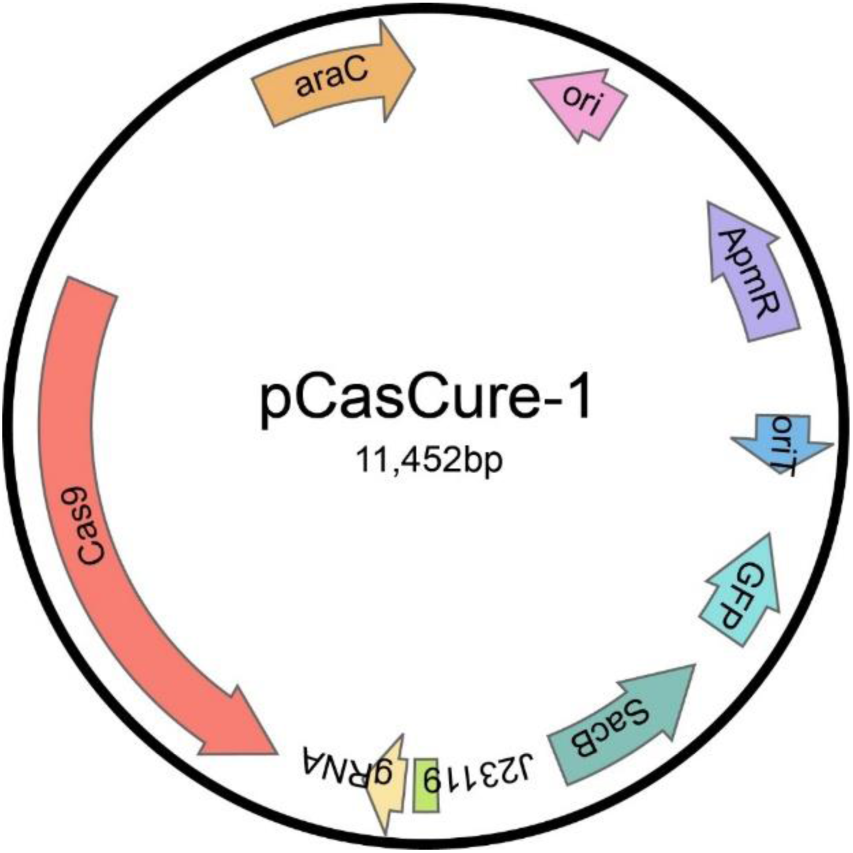
Plasmid map of pCasCure-1. Coding genes are indicated by colored arrows. oriT, the origin of transfer; ori, ColE1 origin of replication; ApmR, apramycin resistance cassette; araC, L-arabinose regulatory protein.

**Figure S2.**
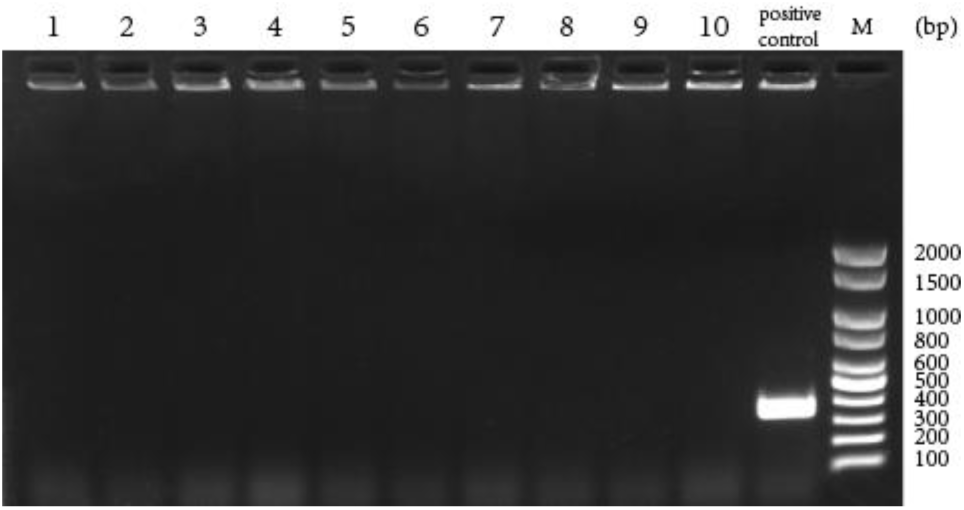
Validation of CRISPR/Cas9-Mediated Curing of the Virulence Plasmid in Klebsiella pneumoniae JNQH1480. Ten colonies were randomly selected following CRISPR/Cas9-based plasmid curing and screened for the presence of the IncHI1B replicon. All colonies (10/10) tested negative, confirming successful elimination of the virulence plasmid. The wild-type strain JNQH1480 served as the positive control. Lane M indicates the DNA size marker; band sizes are annotated adjacent to the gel image.

**Figure S3.**
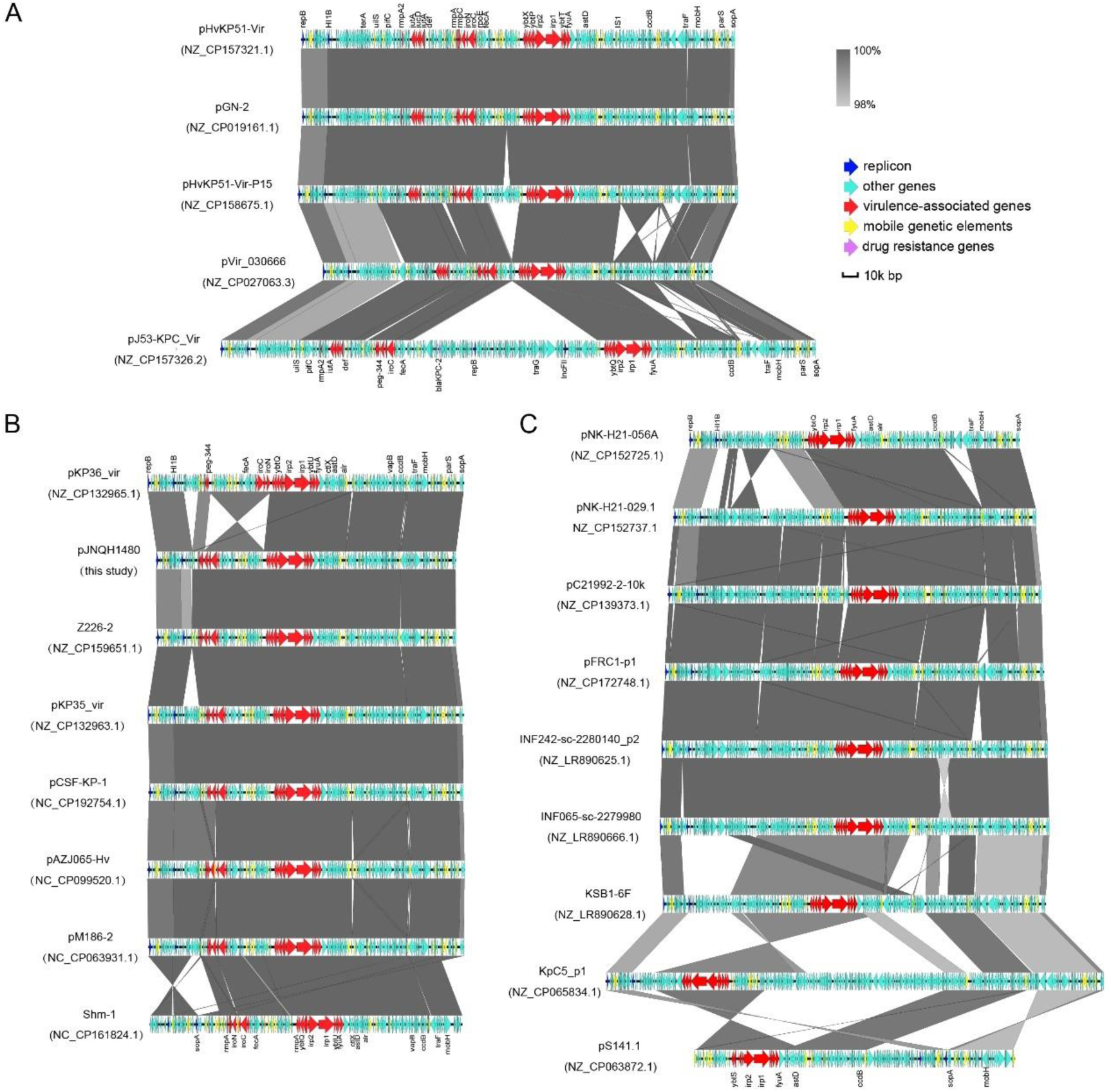
Comparative organization of KpVP-3–like plasmid patterns. Pattern 1 plasmids encode the *ybt* cluster with *irp* and *fyuA* (A). Pattern 2 additionally include *iro*, *rmpADC*, and *peg344* (B). Pattern 3 carry the full virulence repertoire (*rmpADC*, *rmpA2D2*, *iuc*, *iut*, *peg344*, *iro*, *ybt*, *irp*, *fyuA*) (C). Arrows indicate functional categories: red (virulence), yellow (mobile elements), dark blue (replicons), pink (resistance) and light blue (other regions).

---

**Table 1.**
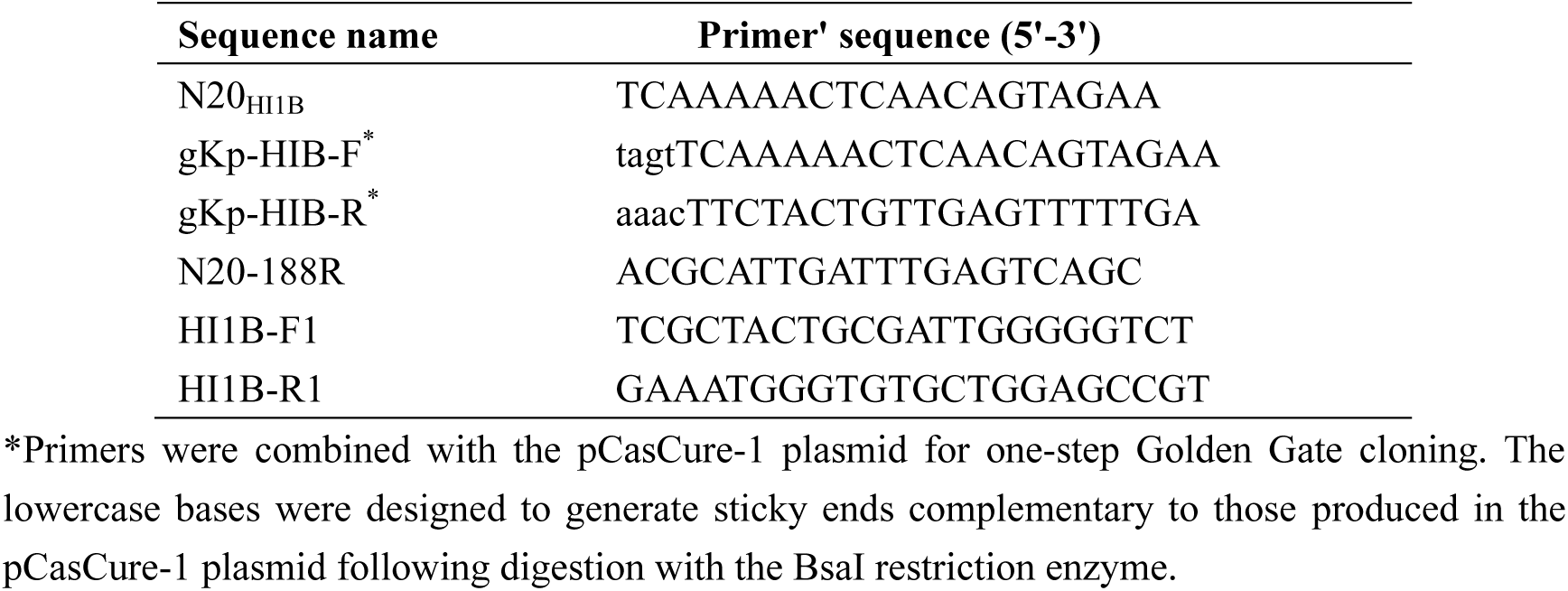
Primers and plasmids used in this study.

**Table S1.** MIC profiles of the hvKp strains isolated in this study and the pVir cured mutant (△pVir) (μg/ml)

| Isolates | β-lactams | | | | | | | | | | Aminoglycosides | | | Quinolones | | others | |
| --- | --- | --- | --- | --- | --- | --- | --- | --- | --- | --- | --- | --- | --- | --- | --- | --- | --- |
|  | AMP | TZP | CZO | CXM | FEP | ATM | IPM | MEM | CAZ | CRO | AMK | TOB | GEN | CIP | LEV | SXT | COL |
| KP1 | >32 | <4 | <4 | <1 | <1 | <1 | <1 | <0.25 | <1 | <1 | <2 | <1 | <1 | <0.25 | <0.25 | <20 | 1 |
| KP2 (JNQH1480) | >32 | <4 | <4 | <1 | <1 | <1 | <1 | <0.25 | <1 | <1 | <2 | <1 | <1 | <0.25 | <0.25 | <20 | 1 |
| KP3 | >32 | <4 | <4 | <1 | <1 | <1 | <1 | <0.25 | <1 | <1 | <2 | <1 | <1 | <0.25 | <0.25 | <20 | 1 |
| KP4 | >32 | <4 | <4 | <1 | <1 | <1 | <1 | <0.25 | <1 | <1 | <2 | <1 | <1 | <0.25 | <0.25 | <20 | 1 |
| KP5 | >32 | <4 | <4 | <1 | <1 | <1 | <1 | <0.25 | <1 | <1 | <2 | <1 | <1 | <0.25 | <0.25 | <20 | 1 |
| JNQH1480ΔpVir | >32 | <4 | <4 | <1 | <1 | <1 | <1 | <0.25 | <1 | <1 | <2 | <1 | <1 | <0.25 | <0.25 | <20 | 0.5 |

AMP, ampicillin; TZP, piperacillin–tazobactam; CZO, cefazolin; CXM, cefuroxime; FEP, cefepime; ATM, aztreonam; IPM, imipenem; MEM, meropenem; CAZ, ceftazidime; CRO, ceftriaxone; AMK, amikacin; TOB, tobramycin; GEN, gentamicin; CIP, ciprofloxacin; LEV, levofloxacin; SXT, trimethoprim–sulfamethoxazole; COL, colistin. All tests were performed in duplicate, and each test included three biological replicates.

**Table S2.**
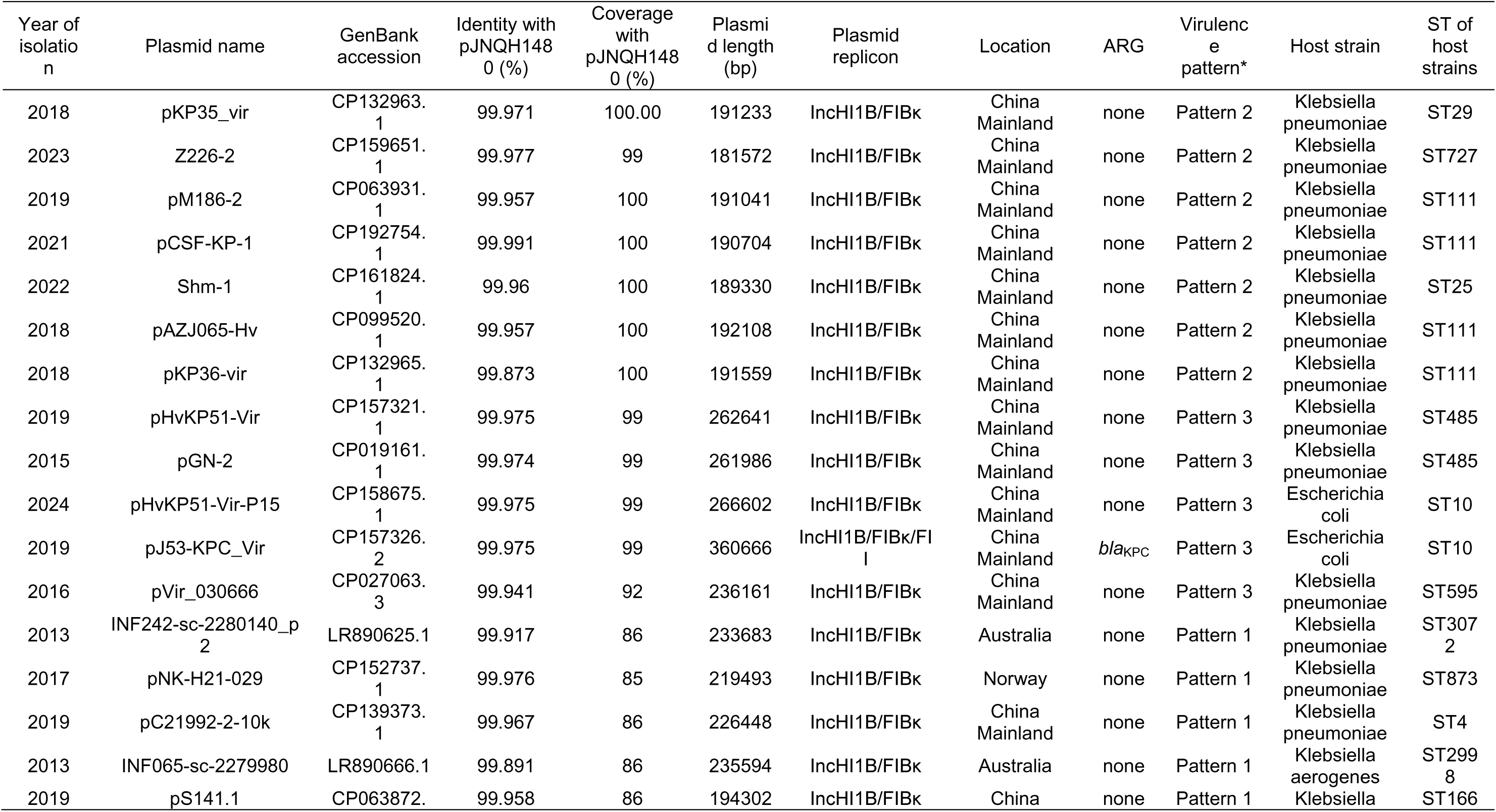
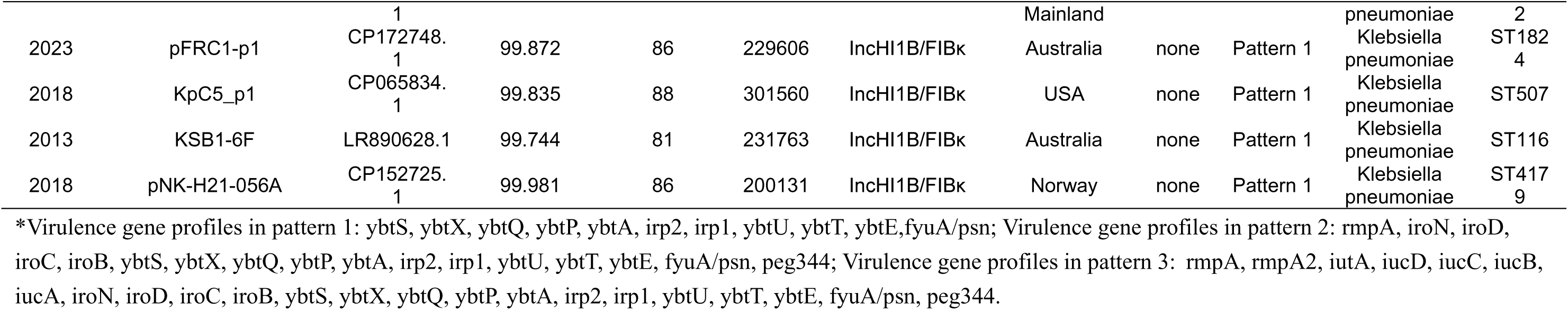
Comparison of virulence plasmid in ST111 hypervirulent *Klebsiella pneumoniae*.

